# A critical role for B cells in cancer immune surveillance

**DOI:** 10.1101/2020.09.19.304790

**Authors:** Kavita Rawat, Anita Tewari, Claudia V. Jakubzick

**Author notes:** Correspondence: Claudia Jakubzick, PhD, Geisel School of Medicine at Dartmouth, Department of Microbiology and Immunology, 628W Borwell, One Medical Center Drive, Lebanon, NH 03756.

## Abstract

It is commonly believed that B cells play no role in cancer immune surveillance. However, this conclusion is based on studies using only a single B cell-deficient mouse strain, muMT mice, which does not show increased tumorigenesis compared with wild-type (WT) mice. In this study, we demonstrate a critical role for B cells in anti-tumor immunity and identify the cellular mechanisms that make muMT mice a special case. First, we replicate previous findings using a melanoma model in muMT mice. Then, we show that in muMT mice, B-cell deficiency is compensated for by a significant increase in another potent anti-tumor cell type, Type-1 interferon (IFN I)-producing plasmacytoid DCs (pDCs), resulting in normative anti-tumor responses. Depleting pDCs in muMT mice resulted in a significant increase in tumor size and burden. Conversely, adoptive transfer of antibodies from naïve WT serum into pDC-depleted muMT mice significantly decreased the tumor load to WT levels. Additionally, a B cell antibody repertoire-deficient mouse strain, IghelMD4 mice, showed a 3-fold increase in tumors relative to WT mice. Overall, these findings indicate the need for a diverse antibody repertoire for early neoplastic cell recognition and the critical role B cells play in anti-cancer immunity.

## Introduction

Tumorigenesis occurs when normal cells accumulate mutations and proliferate out of control. These mutations may result in abnormal cells escaping the body’s first line of defense, intrinsic tumor suppressive mechanisms. The second line of defense is extrinsic tumor suppression, also referred to as cancer immunoediting. This defense is led by immune cells and divided into three stages: elimination (i.e., cancer immunosurveillance), equilibrium, and escape ^1,2^. During the elimination phase, the identification and destruction of nascent tumors result from the expression of neoantigens by mutant cells, forming a hypoxic cluster, and releasing danger-associated molecular patterns (DAMPs), these events alert the immune system of cellular abnormalities. To mount an immune response against neoantigens (as with other foreign antigens), dendritic cells need to become activated and licensed to present neoantigens in an immunogenic way to T cells ^3^. DAMPs and pathogen-associated molecular patterns (PAMPs) are mediators shown to license dendritic cells ^4,5^. All in all, mononuclear phagocytes (dendritic cells, macrophages, and monocytes), NK cells, and T cells are immune cells that play a role in cancer immunosurveillance and the elimination of nascent mutant cells ^6^.

We propose that B cells provide another potential pathway for cancer immunosurveillance. Though their role in tumor neoantigen recognition is not well-established, B cell-produced antibodies are critical for recognizing many other types of neoantigen-expressing cells. For example, because females lack Y-chromosome antigens, syngeneic male cells adoptively transferred into female mice are recognized by natural antibodies and rejected ^7^. Cells bound with antibodies are targeted for elimination and induce an adaptive immune response, which in turn causes a cytotoxic T cell response against H-Y antigen-expressing, male cells. Notably, this immune surveillance and response occurs in the absence of PAMPs, DAMPs, and hypoxic conditions, and with normal MHC-I expression. Similarly, this mechanism of neoantigen recognition and rejection is not based on sex-mismatch. 129Sv female cells adoptively transferred into MHC-matched C57BL/6 female mice also require natural antibodies to start the process of immune rejection ^8^, even when mononuclear phagocytes, NK cells and T cells are present ^7,9^. In sum, in these studies, B cells initiate and play a critical role in neoantigen-expressing cell recognition and elimination. It is unclear why natural antibodies would not play a similar role for the recognition of nascent tumor neoantigens.

Currently, B cells are thought to have no role in cancer immune surveillance in large part because a series of widely cited studies found that B-cell deficient muMT mice mount a robust anti-tumor immune response similar to or greater than wild-type (WT) mice (Extended Data Table 1). A stronger anti-tumor immune response in B cell deficient mice would suggest that B cells inhibit, rather than mediate, anti-tumor immunity. However, several converging lines of evidence challenge this view. At least three other, less known strains of B cell-deficient mice show markedly enhanced tumor growth (Extended Data Table 1). In addition, another method for studying the role of B cells, antibody depletion with anti-CD20 or anti-IgM, shows increased tumor growth, contradicting the muMT findings and suggesting that B cells may mediate anti-tumor immunity (see, Extended Data Table 1). These discrepancies led us to hypothesize that the muMT mice possess compensatory immune mechanisms that prevent tumor growth in the absence of B-cells. If so, then an important role of B-cells in antibody-mediated tumor suppression is masked in the muMT mouse strain.

## Results

### muMT mice have excess pDCs located in the absentee B cell zone

We first assessed tumor burden in the B16F10 melanoma model in two mouse strains with B cell-related deficiencies—B cell-deficient muMT mice and antibody repertoire-deficient IghelMD4 mice—and WT controls. In IghelMD4 mice, >90% of IgM-secreting B cells are specific for hen egg lysozyme. Thus, they lack a complete antibody repertoire. In a previous study, IghelMD4 mice showed enhanced tumor burden compared with WT mice ^7^. The association with high tumor growth provides evidence that B cells participate in cancer immunosurveillance. We replicated the IghelMD4 mouse findings here (Fig.1a and Extended Data Fig. 1). Moreover, the results from muMT mice also replicated earlier results (Extended Data Table 1) showing normal tumor burden compared to WT mice. As anticipated, the overall tumor counts in the B-cell deficient mouse strain, muMT mice, were similar to (though slightly less than) WT mice (Fig. 1a). Thus, muMT mice mount an anti-tumor response in the absence of B cells.

**Fig. 1.**
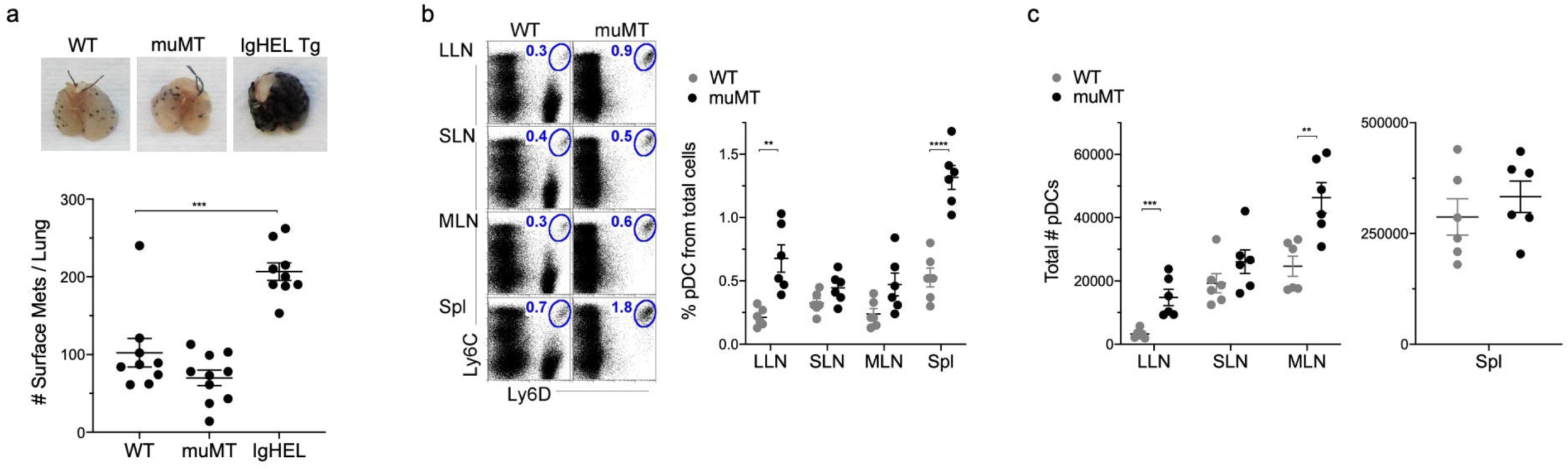
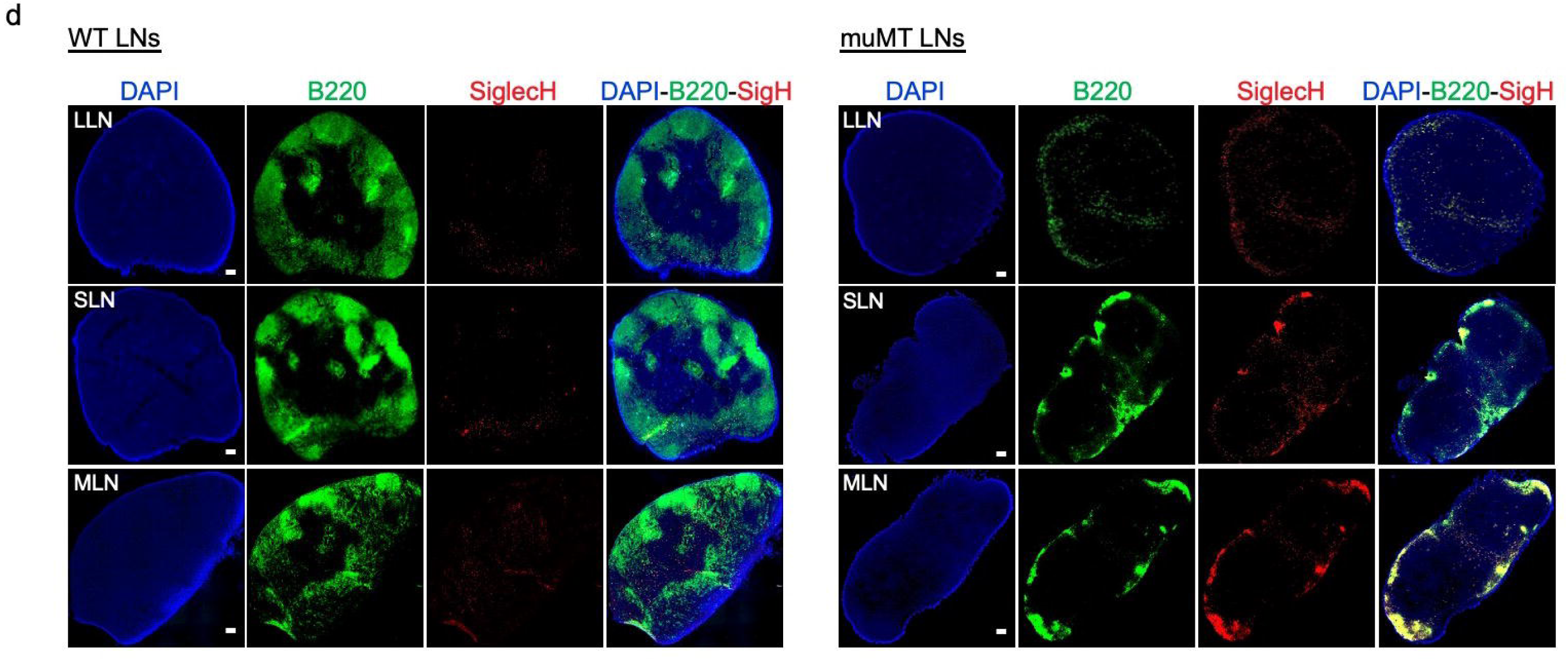
MuMT mice have elevated levels of pDCs in lymph nodes and spleen. (A) WT, muMT and IghelMD4 (IgHEL) mouse lungs were inflated 16 days after i.v. B16F10 challenge. Pics depict total surface metastases (mets) per lung, which were enumerated and illustrated by scatter plot, each dot represents one mouse. Combined data of two independent experiments with 4–5 mice per group. \*\*\**P*<0.0006. (B,C) Left, flow cells plotted as Ly6C versus Ly6D to identify pDCs in lung draining-LNs (LLN), skin draining-LNs (SLN), mesenteric draining-LNs (MLN) and Spleen (Spl). Right, scatter plot displays the pDC frequency and count in WT and muMT mice. Data are representative two combined out of four independent experiments with 3 mice per group. **P<0.0097, ****P<0.0001, (D) Immunohistochemistry (IHC) of LLN, SLN and MLN from naïve WT and muMT mice. Sections were stained with DAPI (blue), anti-B220 (green) and anti-SiglecH (red). Scale bars, 100□μm.

The difference in tumor burden across the two strains, both deficient in B cell-related function, led us to probe more deeply into whether muMT mice show abnormalities in other anti-tumor cell types that could compensate for B cell deficiency, particularly mononuclear phagocytes and pDCs. We observed that the mononuclear phagocyte frequencies were similar between naive muMT and WT mice (data not shown). However, pDCs, lymphoid in origin ^10-13^, were present in significantly greater frequency and numbers in muMT mice compared to WT mice (Fig. 1b-c). We identified them by gating on the intersection of Ly6C and Ly6D expression, and verified that they also express CD11c, B220, and SiglecH and lack the expression of CD11b and CD19 (Extended Data Fig. 2). PDCs are potent anti-viral and anti-tumor cells ^14 15^, this indicates a potential mechanism that compensates for B cell deficiency in muMT mice.

We next characterized the location of pDCs within the LNs. In the paracortex and medulla regions, we observed SiglecH^+^ pDCs in both muMT and WT mice (Fig. 1c and Extended Data Fig. 2). However, we found marked differences in the region adjacent to the subcapsular sinus, with few pDCs in WT mice but concentrated clusters of pDCs in muMT mice (Fig. 1c and Extended Data Fig. 3a). This region is termed the ‘B cell zone’ in WT mice, and we refer to this as the ‘absentee B cell zone’ in muMT mice. These findings suggest that muMT mice have increased levels of pDCs where B cells would normally reside within draining LNs, providing further evidence for a compensatory mechanism in muMT mice.

### In the absence of B cells, a significant number of IFN I-producing pDCs accumulate in tumors in muMT mice

We next turned to the melanoma model, examining whether muMT mice show increased pDC counts and function in the lung tumor environment compared to WT mice. We analyzed Type 1 interferon (IFN 1), IFN-alpha, a robust anti-tumor cytokine that is produced by activated pDCs in larger quantities than any other immune cell type ^16-18^. PDC counts were increased by 4.6-fold in the tumors of muMT mice, along with elevated levels of IFN-alpha, compared with WT mice (Fig. 2a and Extended Data Fig. 3b). Elevated expression of IFN I-alpha was also observed in IHCs of the lung draining-LNs of tumor-bearing muMT mice compared to WT mice (Fig. 2b). Thus, an increased influx of IFN I-producing pDCs in the tumor may explain why muMT mice do not show enhanced tumor burden in the absence of B cells.

**Fig. 2.**
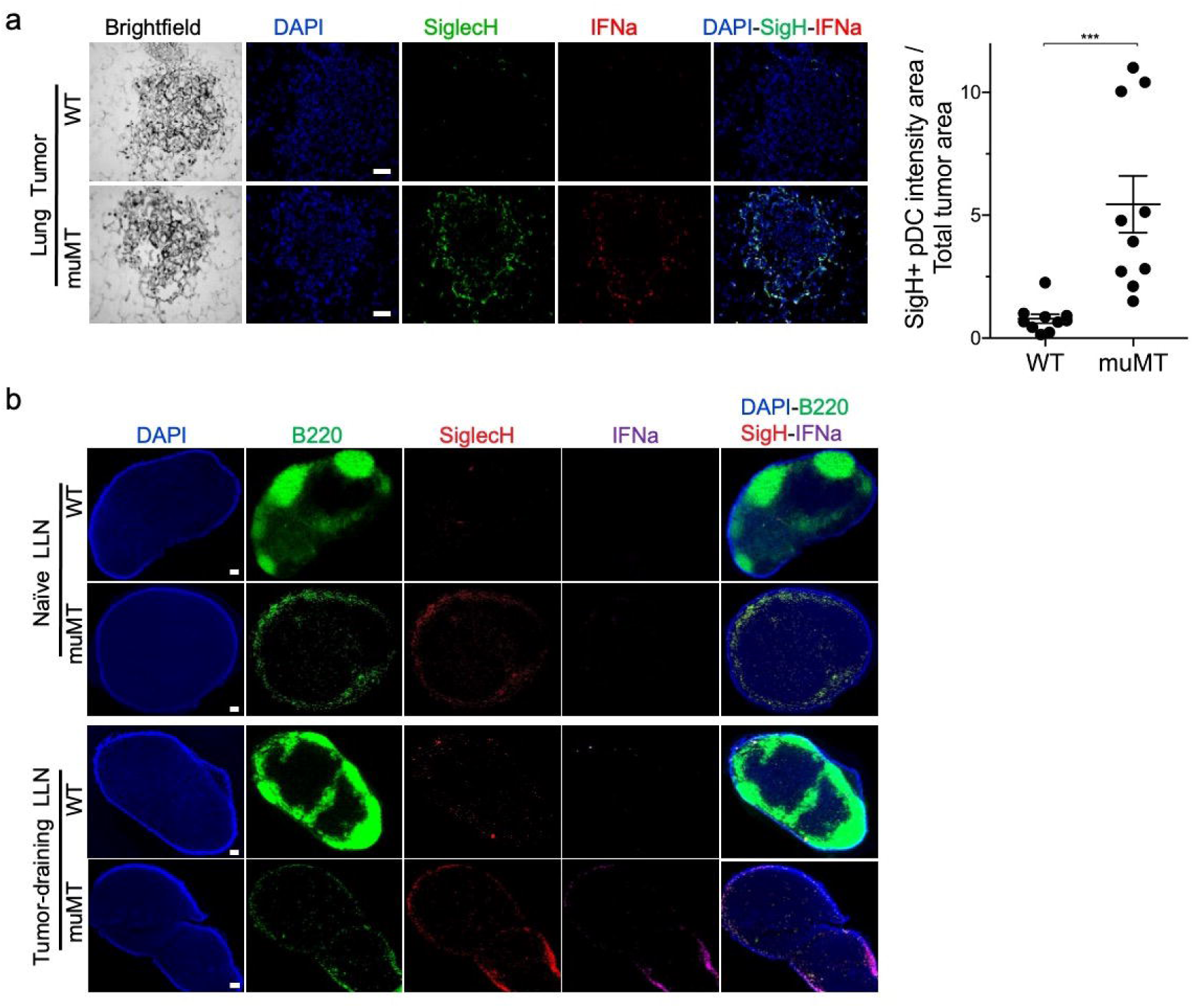
Enhanced accumulation of IFN-a secreting pDCs at the lung melanoma site was observed in muMT mice compared to WT. (A) IHC of lung melanoma in WT and muMT mice stained with DAPI (blue), anti-SiglecH (green) and anti-IFNa (red). Scale bars, 40_μm. Scatter plot illustrates Kenyence quantitative software analysis of SigH+ intensity over total tumor area X 100. Each dot plot represents one tumor of two tumors per mouse from 5 mice per group. Data are representative of three independent experiments. (B) IHCs of LLNs from naïve and melanoma induced WT and muMT mice. Sections were stained with DAPI (blue), anti-B220 (green), anti-SiglecH (red) and anti-IFNa (purple). Scale bars, 100□μm.

### PDC depletion in muMT mice increases tumor burden

To verify the functional role of pDCs in tumor suppression, we selectively depleted pDCs with targeted antibodies against SiglecH and B220 (see Supplementary Materials for details). SiglecH is a selective pDC marker, and in muMT mice, all B220^+^ cells in LN and tumors are SiglecH^+^ (Extended Data Fig. 1-3). PDC-depleted muMT mice showed a 3-fold increase in tumor counts, along with tumor size, compared with both non-depleted muMT and WT mice (Fig. 3a and Extended Data Fig. 5a). We hypothesized that in the absence of the compensatory pDC mechanism, natural antibodies are essential for neoantigen recognition on the surface of cancer cells. To test this, pDC-depleted muMT mice were injected with melanoma and antibody-rich naive WT serum. Antibody restoration resulted in a significant 2.2-fold decrease in lung tumors compared to pDC-depleted muMT mice (Fig. 3b and Extended Data Fig. 5b). Thus, natural antibodies appear to be essential for the recognition and elimination of melanoma tumor cells in muMT mice when the compensatory pDC mechanism is blocked.

**Fig. 3.**
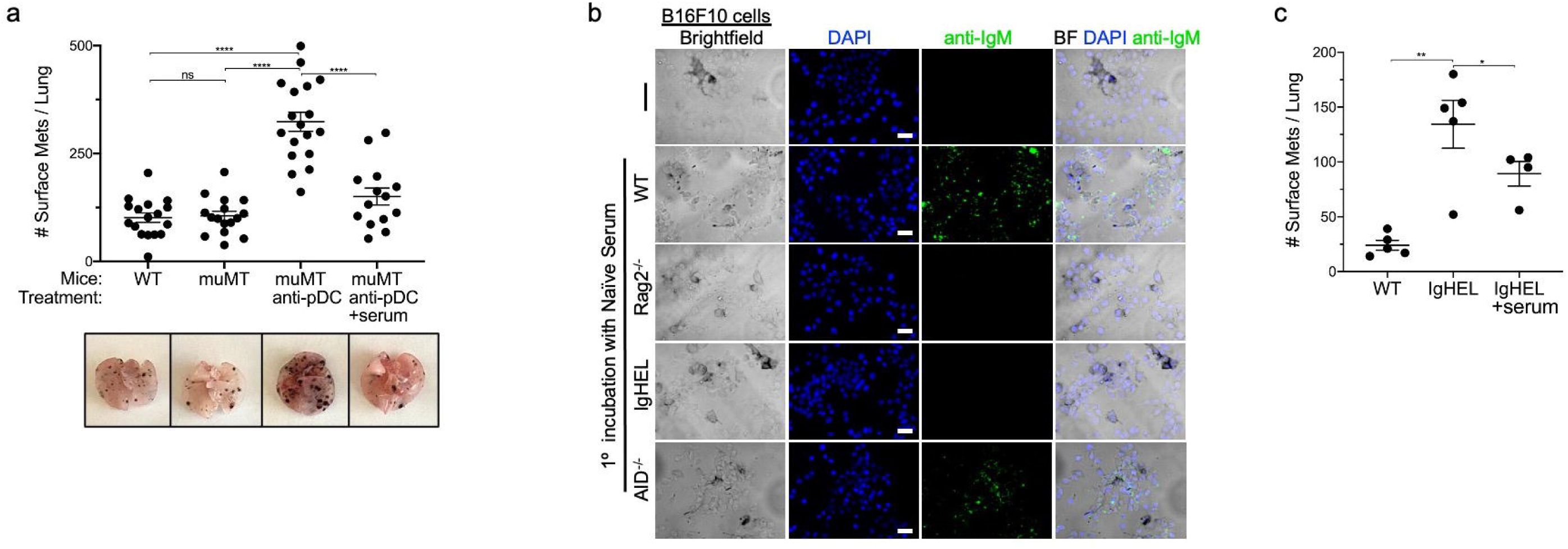
PDC depletion in muMT mice results in increased tumor burden, highlighting the importance of B cells and antibody repertoire in cancer immune surveillance. (A) WT and muMT mouse lungs were inflated 16 days after i.v. B16F10 melanoma cell lines. Pics depict total surface metastases (mets) per lung, which were enumerated and illustrated by scatter plot, each dot represents one mouse. Two muMT mouse groups were pDC-depleted using a triple Ab treatment every 48 hours up to harvest day. Combined data of three independent experiments with 4–5 mice per group. ****P <0.0001. Non-significant (ns). Mean ±SEM. (B) Lab-Tek Chamber Slides were plated with live B16F10 cells and first incubated for 30 mins with either naïve WT, Rag2^-/-^, IghelMD4, and AID^-/-^ serum or no serum. Then, cells were washed and incubated with secondary mouse anti-IgM (green). Scale bars, 40□μm. (C) WT and IghelMD4 mouse lungs were inflated 16 days after i.v. B16F10 melanoma cell lines and enumerated and illustrated by scatter plot, each dot represents one mouse. Two muMT groups were treated with pDC depleting triple Ab every 48 hours up to harvest day. Combined data of three independent experiments with 4–5 mice per group. ****P <0.0001. Non-significant (ns). Mean ±SEM.

To directly demonstrate that antibodies in naive WT serum recognize live melanoma cells, we cultured B16F10 melanoma cells on Lab-tek microscope chamber slides and incubated for 30 mins with four different types of naïve serum: (1) WT, rich in natural IgM and other antibodies; (2) IghelMD4, which lack a complete antibody repertoire, with antibodies IgM-specific only for the HEL epitope; (3) Rag2^-/-^, which contain no antibodies; and (4) AID^-/-^, which have a hyper-IgM repertoire because they lack the ability to isotype switch [no IgG] and perform somatic hypermutation, but is otherwise complete natural IgM antibodies. If a complete IgM antibody repertoire is important for cancer immunosurveillance, the WT and AID^-/-^ serum groups alone should show binding of antibodies on live tumors. This was the case. Binding of natural IgM antibodies was robustly detected on melanoma cells treated with naïve WT and AID^-/-^ serum, but not Rag^-/-^ or IghelMD4 serum (Fig. 3b).

Finally, returning to in vivo studies, we compared the tumor burden in two groups of IghelMD4 mice: One treated with naive WT serum and one untreated. Antibody enrichment with WT serum significantly decreased tumor counts compared with untreated IghelMD4 mice (Fig. 3c). All in all, these data support the hypothesis that B cell secreting natural antibodies are critical for detection and elimination of nascent cancer cells—unless abnormal compensatory mechanisms develop, such as the increased tumor infiltration of IFN-I producing pDCs.

## Discussion

In this study, we show a critical role for B cells and natural antibodies in cancer immunosurveillance using a melanoma model. The findings here was facilitated in part by the use of IghelMD4 mice to investigate neoantigen recognition, rather than the more commonly used muMT strain. We used IghelMD4 mice in a prior study because muMT mice lack appropriate lymphatic development in the LN cortex due to the absence of B cells ^19^. We wanted to study cancer immunosurveillance in a model with normative lymphatics. This led us down the road of comparing different types of B cell, antibody-deficient mice.

We set out to understand why muMT mice develop a robust early anti-tumor immune response in the absence of B cells and natural antibodies, particularly when antibodies are critical for other forms of neoantigen recognition and other B cell or antibody-deficient mouse strains show increased tumor proliferation (Extended Data Table 1). We replicated previous findings that antibody-deficient IghelMD4 mice displayed significantly more tumors than WT mice ^7^, and muMT mice do not. We then show that negative findings in muMT mice are due to a compensatory pDC mechanism. In naïve muMT mice, pDCs are increased in the absentee B cell zone of the lymph nodes. After melanoma injection, pDCs were found to infiltrate tumors and express IFN 1, a potent tumor-suppressing cytokine. PDC-deficient muMT mice, in whom this compensatory mechanism is disrupted, show dramatically increased tumor burden, but serum injections that restore the complete antibody repertoire are sufficient to restore anti-tumor immunity.

This study does not challenge the empirical validity of previous findings in muMT mice, which are reproducible (Extended Data Table 1). Instead, it highlights that relying on results from a single mouse strain can lead to erroneous conclusions. MuMT mice differ from WT in their lack of B cells, but also in other respects, including increases in tumor-suppressive pDC cells; thus, the assumption that using B cell-deficient mice provides a pure test of B cell function is, in this case, unwarranted. Converging evidence from other B cell-deficient and antibody-deficient strains, antibody depletion, and antibody restoration paints a different and more complete picture: Antibodies produced by B cells are a critical part of the cancer immunosurveillance repertoire, but in specific cases, including muMT mice, other systems can compensate, masking the importance of B cells in these specific models.

Our findings raise the question of why B cell deficient muMT mice exhibit increased pDC frequencies. pDCs and B cells share a common lymphoid progenitor ^10,11^, and perhaps the inability of the progenitor to become a B cell results in the differentiation of pDCs. As pDCs normally represent a small fraction of immune cells in the lymphoid compartment, they may not be present in sufficient numbers to initially enter in vast quantities in the tumor environment in WT mice, as opposed to what we observed in muMT mice. Another interesting observation was the clustering of pDCs in the absentee B cell zone of muMT mice. This was not observed in WT mice (Fig. 1c). The implications of pDCs residing in the B cells zone of the LNs is unclear, perhaps it is a site for pDC activation.

Future studies could productively explore the mechanisms by which antibodies recognize and tag tumor cells, which could be exploited in treatment development. Natural antibodies have a low affinity for self-antigens ^20 21,22^, and may increase their affinity if a given cell surface antigen mutates or alters its glycosylation. It is known that antibodies on the surface of a cell tags it for elimination by recruiting complement components and innate immune cells, and can promote the activation of adaptive immune cells by presenting altered self-antigens in an immunogenic fashion ^7^. In conclusion, these findings challenge the long-standing view that B-cells are not important for cancer immunosurveillance, and show instead that they play a vital role in anti-tumor immunity.

## Methods

### Mice

CD45.2 wild-type (WT), B6.129S2-*Ighm*^*tm1Cgn*^/J (muMt), AID^-/-^ and IghelMD4 mice were purchased from Jackson Research Laboratories and Charles River NCI. All mice were bred in house. Mice were genotyped or phenotyped prior to studies and used at 6–8 weeks of age, housed in a specific pathogen-free environment at Dartmouth Hitchcock Medical College, an AAALAC accredited institution, and used in accordance with protocols approved by the Institutional Animal Care and Utilization Committee.

### Flow Cytometry

Tissues were minced and digested with 2.5□mg/ml collagenase D (Roche) for 30□min at 37□°C. 100□μl of 100□mM EDTA was added to stop 1□ml of enzymatic digestion. Digested tissue was pipetted up and down 30 times using a glass Pasteur pipette and passed through a 70-μm nylon filter to acquire single-cell suspensions from lungs, spleen, lung draining-LNs, skin draining-LNs, and mesenteric draining-LNs. Cells were stained with the following monoclonal Abs: Phycoerythrin (PE)-conjugated to Siglec H, Ly6G and NK1.1; PerCP-Cy5.5-conjugated to B220; PE-Cy7-conjugated to CD11c and CD44; BUV395-conjugated to CD11b and CD4; fluorescein isothiocyanate (FITC)-conjugated to Ly6D, CD3 and CD62L; allophycocyanin-conjugated to CD19 and CD103; APC-Cy7-conjugated to CD45; and BV510 conjugated to Ly6C. The viability dye DAPI (#D9542, Sigma) was added immediately before each sample acquisition on a BD Symphony A3 analyzer (BD Biosciences). Data were analyzed using FlowJo (Tree Star, Ashland, OR). Antigen-specific antibodies and isotype controls were obtained from BioLegend, eBioscience and BD Biosciences.

### B16F10 lung melanoma model

B16F10 melanoma cells were purchased from ATCC (CRL-6475) and maintained in RPMI with 10% FCS, 1% Pen/Strep/L-glutamine (Sigma), 1% non-essential amino acids (Sigma), 1% sodium pyruvate (Sigma), 10□mM HEPES (Sigma) and 0.1□mM β-mercaptoethanol. Mice were intravenously challenged with 2 × 10^5^ viable B16F10 cells and euthanized 16 days post injection. Lungs of mice were inflated with 1% agarose. A blinded observer counted the B16F10 lung surface metastases.

### Plasmacytoid dendritic cells (pDC) depletion in muMT mice

To deplete pDCs in muMT mice, 1 mg of anti-Siglec H (clone 440c Rat IgG2b, BioXcell), 1mg anti-mouse B220, (clone RA3.3A1/6.1 Rat IgM, BioXcell) and 1 mg anti-B220, (clone RA3-6B2 Rat IgG2a, Leinco technologies, Inc) were injected individually and in combination into the peritoneal cavity of the mice. Assessment of pDC depletion from the spleen, LLN, SLN and MLN was performed at 24, 48 and 72 hours. To deplete pDCs long-term in the B16F10 melanoma model, mice were i.p. injected with the triple antibody (Ab) treatment every 48 hours until harvest, day 16. A group of pDC-depleted mice were also i.p injected with 200 μl of native WT serum every 48 hours until lung harvest.

### Microscopy

Lungs and LNs were excised and immersed in 4% paraformaldehyde with 10% sucrose and 7% picric acid in PBS for 2 hours and then embedded in Tissue-Tec OCT (Fisher Scientific). 10 micron sections were prepared and slides were stained for B220, SiglecH and IFNa. Control slides were treated with Ab isotype controls and secondary antibodies. Stained slides were mounted with Prolong Gold Antifade containing DAPI (# P36931, Invitrogen), whole lymph node images were stitched using 10X objective lens and imaged with a Keyence fluorescence microscope (BZ-X800 series).

### Statistics

Statistical analysis was conducted using InStat and Prism software (GraphPad). All results are expressed as the mean ±SEM. Statistical tests were performed using two-tailed Student’s *t*-test. A value of *P*<0.05 was considered statistically significant.

## Acknowledgement

This work was supported by National Institutes of Health grants R01 HL115334 and R01 HL135001 (CVJ).

## Author Contributions

CVJ and KR prepared the manuscript; KR, AT, and CVJ. executed the experiments; all authors provided intellectual input, critical feedback, discussed results, and designed experiments.

## Additional Information

Supplementary Information is available for this paper.

Correspondence and requests for materials should be addressed to Claudia Jakubzick.

## Supplementary Data

**Extended Data Table 1.** Articles not supporting and supporting the hypothesis that B cells are essential for early cancer cell recognition and anti-tumor immunity. *As of August 2020

| Not supporting | Title | Author (ref) | Journal | Year | Mouse model | Cancer type | *No. of Citations |
| --- | --- | --- | --- | --- | --- | --- | --- |
| 1 | The role of B cells for in vivo T cell responses to a Friend virus-induced leukemia. | Schultz et al.<sup>1</sup> | Science | 1990 | muMT | Friend virus-induced leukemia | 99 |
| 2 | B cells inhibit induction of T cell-dependent tumor immunity | Qin et al.<sup>2</sup> | Nature medicine | 1998 | muMT | TS/A cells, J558L cells | 411 |
| 3 | Combination Chemotherapy and IL-15 Administration Induce Permanent Tumor Regression in a Mouse Lung Tumor Model: NK and T Cell-Mediated Effects Antagonized by B Cells | Chapoval et al.<sup>3</sup> | J Immunol | 1998 | muMT | 76-9 cells (syngeneic B6 3-methylcholanthrene-induced) | 94 |
| 4 | Enhanced efficacy of melanoma vaccines in the absence of B lymphocytes | Perricone et al.<sup>4</sup> | J Immunother | 2004 | muMT | B16F10 | 80 |
| 5 | Increased rejection of primary tumors in mice lacking B cells: Inhibition of anti-tumor CTL and TH1 cytokine responses by B cells | Shah et al.<sup>5</sup> | Tumor Immunology | 2005 | muMT | EL4 thymoma, B16F10 and MC38 colon carcinoma | 197 |
| 6 | Inhibitory Effects of B Cells on Antitumor Immunity | Inoue et al.<sup>6</sup> | Cancer Res | 2006 | muMT | Friend murine leukemia virus gag-expressing mouse EL-4, D5 mouse melanoma, or MCA304 mouse sarcoma cells. | 285 |
| 7 | The absence of B lymphocytes reduces the number and function of T-regulatory cells and enhances the anti-tumor response in a murine tumor model | Tadmor et al.<sup>7</sup> | Cancer Immunol Immunother | 2011 | muMT | EMT-6 mammary adenocarcinoma cells | 100 |
| 8 | Immune-promoted tumor cell invasion and Metastasis New consideration in Cancer Therapy | Guillem and Sampsel<sup>8</sup> | Cancer therapy (Google books) | 2012 | muMT | B16F10 and LLC | 4 |
| 9 | B lymphocyte inhibition of anti-tumor response depends on expansion of Treg but is independent of B-cell IL-10 secretion | Zhang et al.<sup>9</sup> | Cancer Immunol Immunother | 2013 | muMT | EMT-6 mammary tumors | 65 |
| 10 | Allogeneic cancer cell-based immunotherapy | Eckhard R Podack<sup>10</sup> | (US PATENT US8475785B2) | 2013 | muMT | EG7, EL4 and LLC cell line | 5 |
| 11 | B cells promote tumor progression in a mouse model of HPV-mediated cervical cancer | Tang et al.<sup>11</sup> | Int. J. Cancer | 2016 | muMT | TC-1 tumor cells and B16 Ova | 19 |
| 12 | B lymphocytes can be activated to act as antigen presenting cells to promote anti-tumor responses | Rossetti et al.<sup>12</sup> | PLOS ONE | 2018 | muMT | TC-1 cell line | 10 |
| 13 | IL-33 delays metastatic peritoneal cancer progression inducing an allergic microenvironment | Perales-Puchalt et al.<sup>13</sup> | Oncolimmunology | 2018 | muMT | ID8 cell line | 3 |
| 14 | Polynucleotides for inhibiting metastasis and tumor cell growth | Barbera-Guillem<sup>14</sup> | (US PATENT US6268350B1) |  | muMT | B16F10 |  |
| 15 | B cells Regulate Macrophage Phenotype and Response to Chemotherapy in Squamous Carcinomas | Affara et al.<sup>15</sup> | Cancer Cell | 2015 | JH-/- | SCC cell lines derived from K14-HPV16 mice | 169 |
| Supporting | Title | Author (ref) | Journal | Year | Mouse model | Cancer type | *No. of Citations |
| 1 | Anti-tumor immunity in B-lymphocyte-deprived mice. III. Immunity to primary Moloney sarcoma virus-induced tumors | Gordon et al.<sup>16</sup> | Int J Cancer | 1982 | anti-mouse IgM treatment | Moloney sarcoma virus (MSV) induced tumor | 22 |
| 2 | Low natural antibody and low in vivo tumor resistance, in xid-bearing B-cell deficient mice | Chow and Bennet<sup>17</sup> | The Journal of Immunology | 1989 | X-linked immunodeficiency-bearing B-cell deficient mice-low serum IgM and IgG3 | RI-28, a radiation-induced T cell leukemia | 24 |
| 3 | B Cells Are Required for Optimal CD4+ and CD8+ T Cell Tumor Immunity: Therapeutic B Cell Depletion Enhances B16 Melanoma Growth in Mice | DiLillo et al.<sup>18</sup> | J Immunol | 2010 | anti-CD20 mAb treatment | B16F1 | 229 |
| 4 | B cells promote tumor immunity against B16F10 melanoma | Kobayashi et al.<sup>19</sup> | Am J Pathol | 2014 | B-cell linker protein (BLNK) deficient (BLNK-/-) mice, in which B cell development and function is severely impaired | B16F10 | 25 |
| 5 | Immune Surveillance by Natural IgM Is Required for Early Neoantigen Recognition and Initiation of Adaptive Immunity | Atif et al.<sup>20</sup> | Am J Respir Cell Mol Biol | 2018 | IghelMD4 | B16F10 Urethane-induced tumors | 8 |

