## Supplementary File for "A critical role for B cells in cancer immune surveillance"

**Supplementary Data**

**Extended Data Table 1**| Articles not supporting and supporting the hypothesis that B cells are essential for early cancer cell recognition and anti-tumor immunity. *As of August 2020

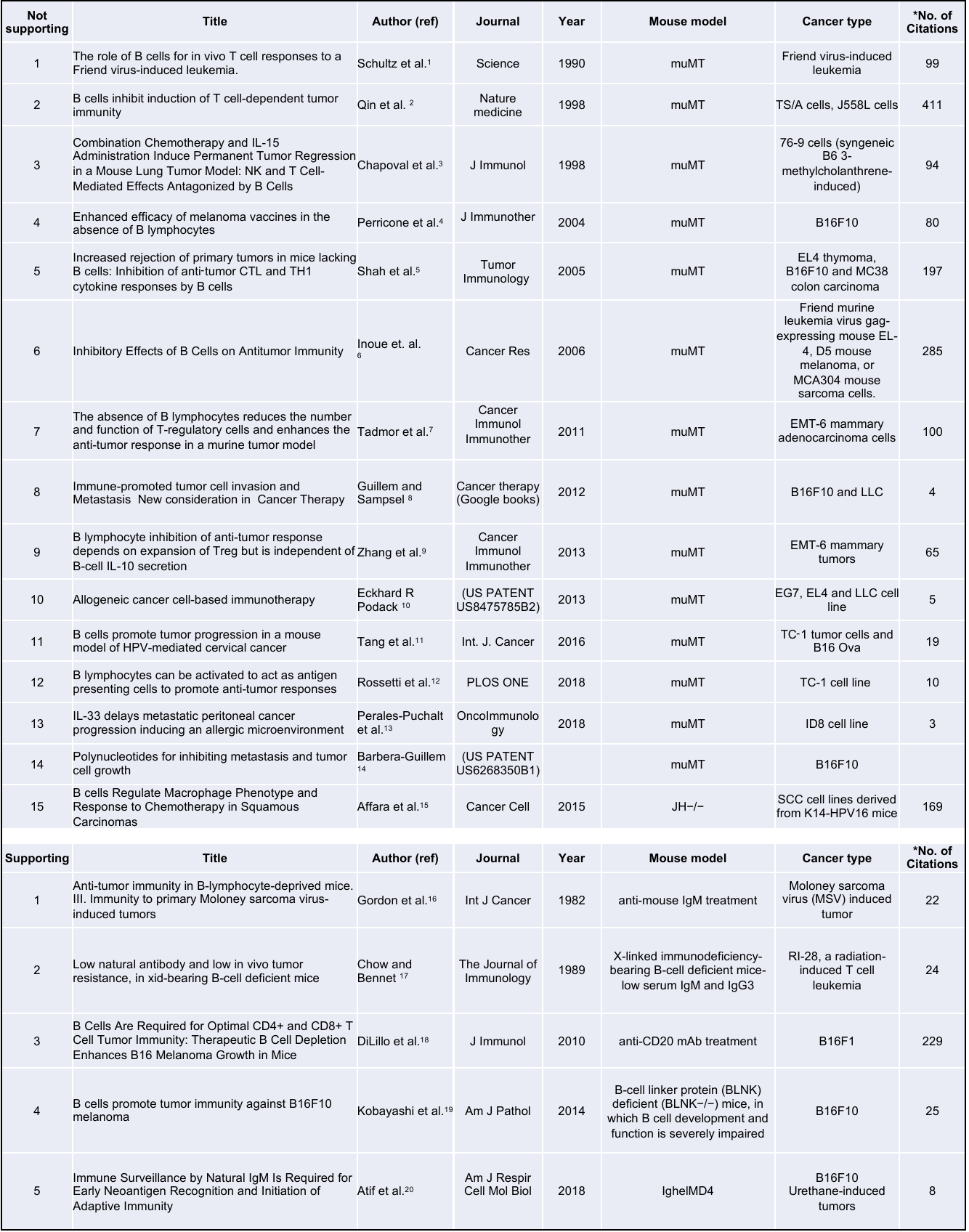

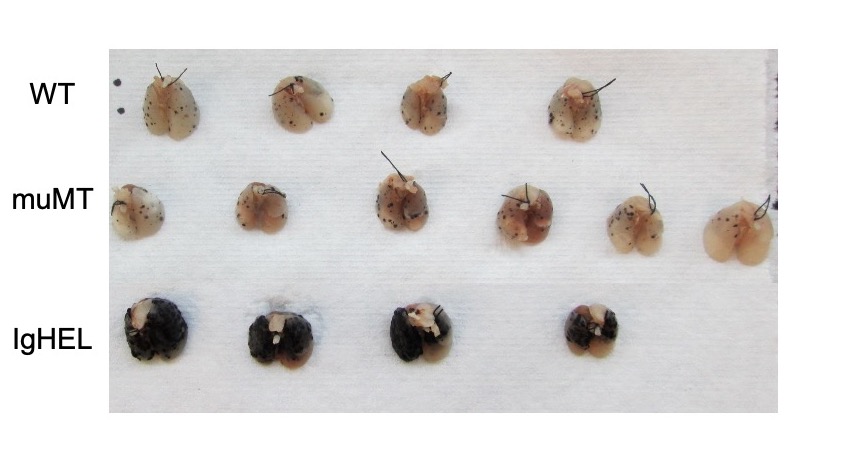

**Extended Data Fig. 1**| **B16F10 melanoma tumor burden in WT, muMT and IghelMD4 mice.** Picture depicts agarose inflated lungs challenged with B16F10 melanoma in WT, muMT, and IghelMD4 mice.

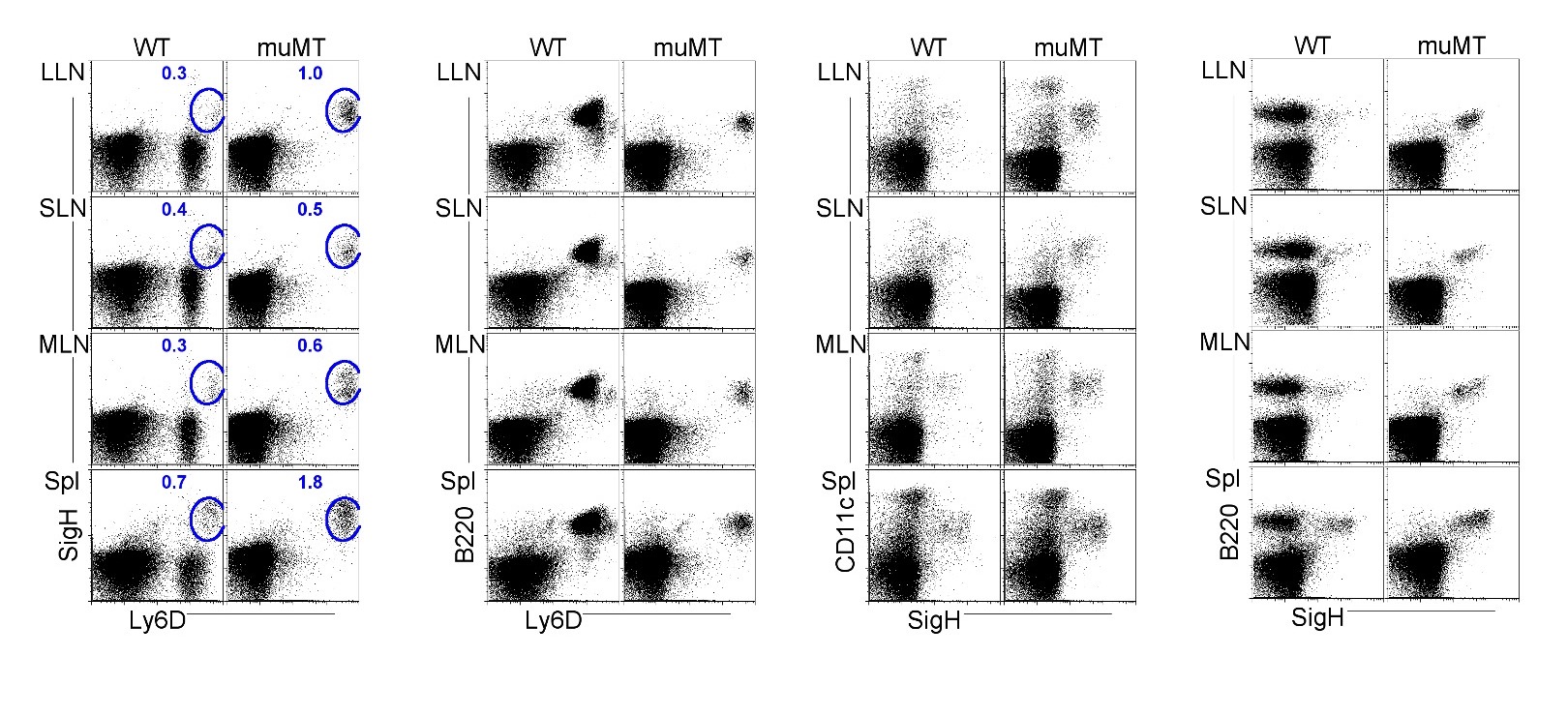

**Extended Data Fig. 2**| **Flow plots illustrate pDC stains and gating strategies.** Flow cells plotted in different ways such as SiglecH versus Ly6D, B220 versus Ly6D, CD11c versus SiglecH, and B220 versus SiglecH to identify pDCs in lung draining-LNs (LLN), skin draining-LNs (SLN), mesenteric draining-LNs (MLN) and Spleen (Spl).

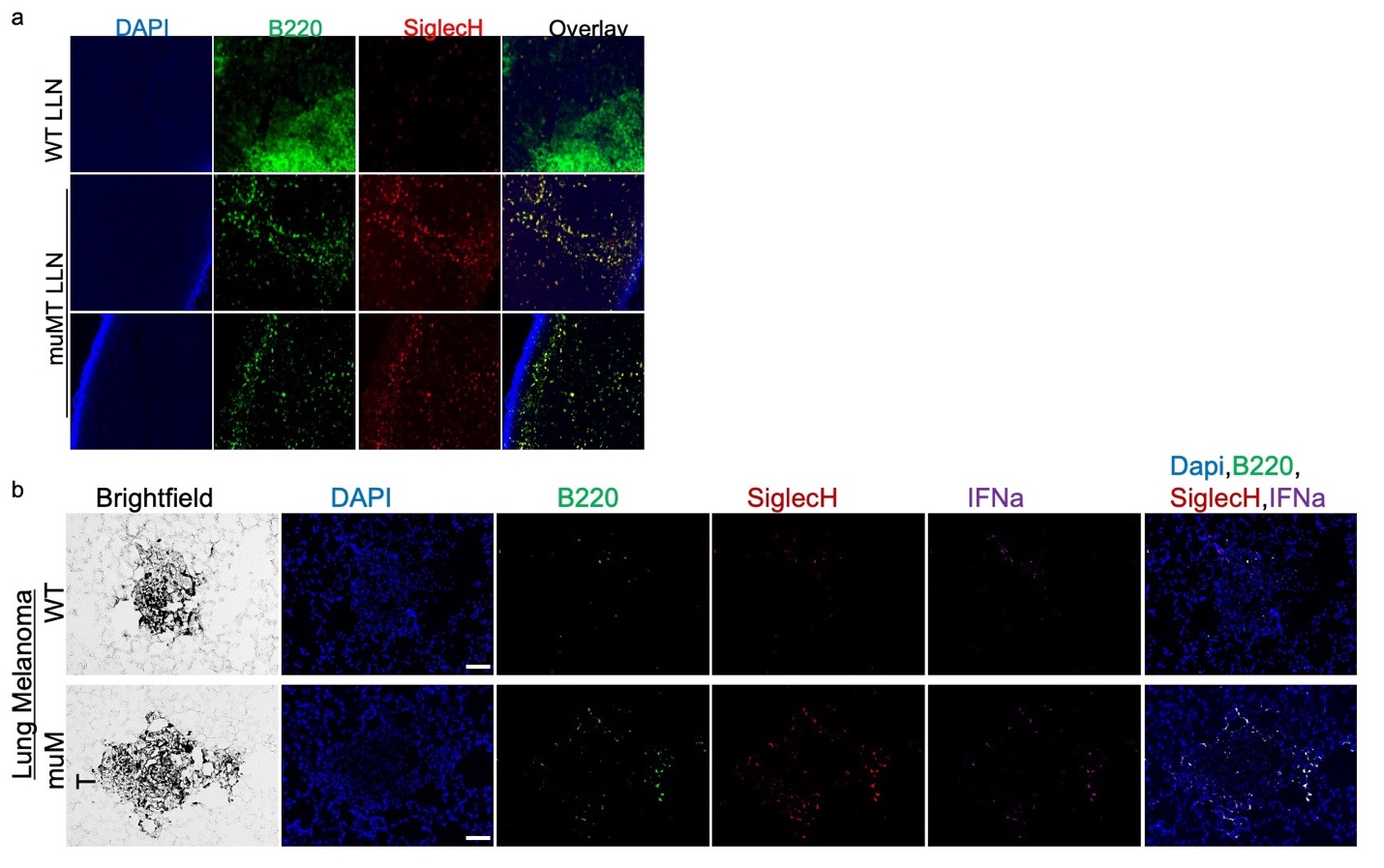

**Extended Data Fig. 3**| **pDC staining of WT and muMT lymph nodes and lung tumors.**

(a) IHC 20X image muMT of LLN and Spl Sections were stained with DAPI (blue), anti-B220 (green) and anti-SiglecH (red). Scale bars, 10 μm

(b) IHC of lung melanoma in WT and muMT mice stained with DAPI (blue), anti-SiglecH (green) and anti-IFNa (red). Scale bars, 40 μm.

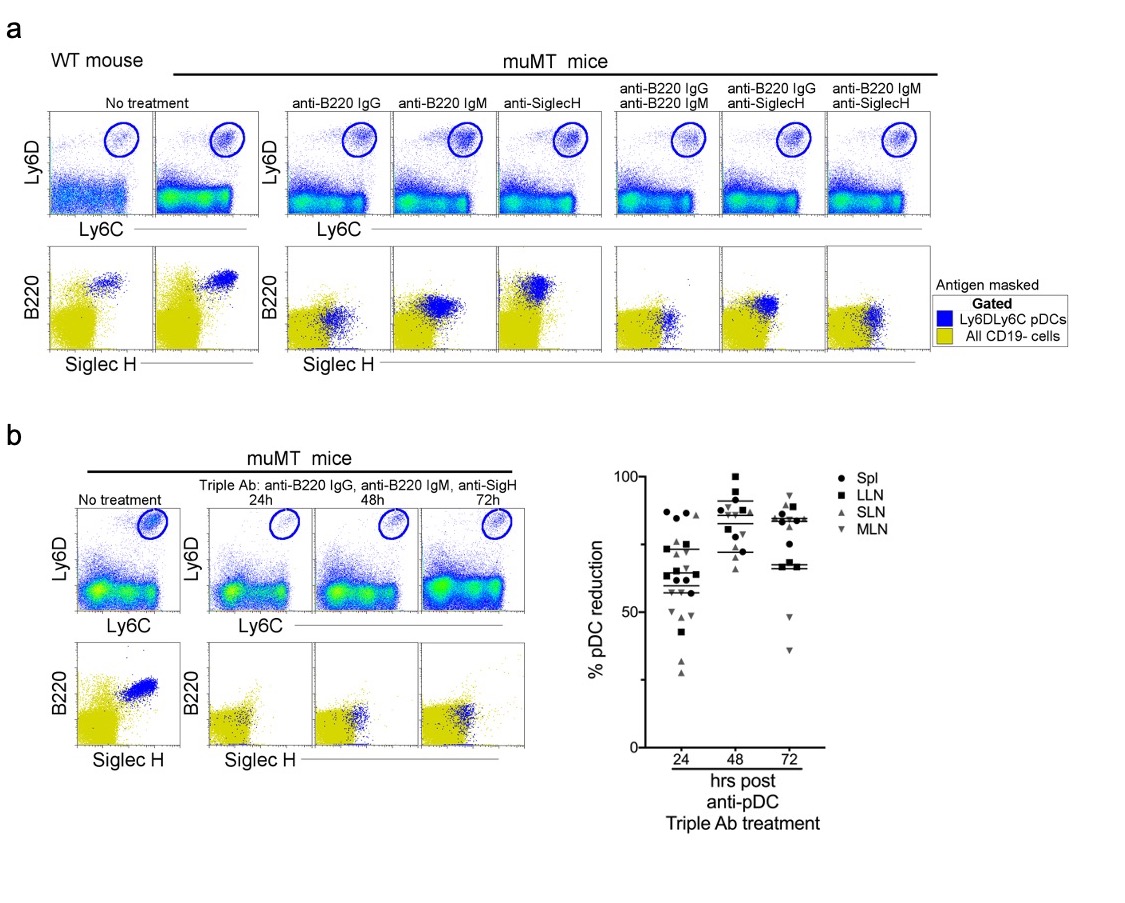

**Extended Data Fig. 4**| **Depletion of the anti-tumor compensatory cell type, pDCs, in muMT mice.** Three distinct monoclonal antibodies were tested for their capacity to deplete pDCs. Two of the antibodies are specific for B220: Rat anti-B220 IgG2a and Rat anti-B220 IgM; and one is specific for Siglec-H: Rat anti-Siglec-H IgG2b. The latter antibody is an isotype shared by other established depleting antibodies such as anti-CD4 (clone GK1.1) and anti-Gr1 (clone RB685C). (a) At high concentrations of 1 mg, none of the antibodies alone, or in combinations of two, depleted pDCs at 24 hours, instead the antibodies masked the targeted antigen as illustrated in flow plot overlays of pDCs identified by Ly6C and Ly6D from all splenic CD19- cells. Bottom row, flow cytometry overlays of all CD19- cells (yellow) and gated pDCs (blue) illustrate the masking of B220 and SiglecH 24 hours after muMT mice were ip injected, alone or in combination, with anti-mouse B220 rat IgG2a, anti-mouse B220 rat IgM, and anti-mouse Siglec-H. (b) However, when all three antibodies were given, pDCs were diminished by 63-83% in the spleen and LNs of muMT mice, and this depletion persisted across 24, 48 and 72 hours**.** Top row, gated Ly6C^+^Ly6D^+^ pDCs from all splenic CD19- cells. Bottom row, flow cytometry overlays of all CD19- cells (yellow) and gated pDCs (blue) illustrating the few cells remaining at 24, 48 and 72 hours after pDC-depleting triple antibody (Ab) treatment (anti-mouse B220 rat IgG2a, anti-mouse B220 rat IgM, and anti-mouse Siglec-H). Scatter plot represents four independent experiments.

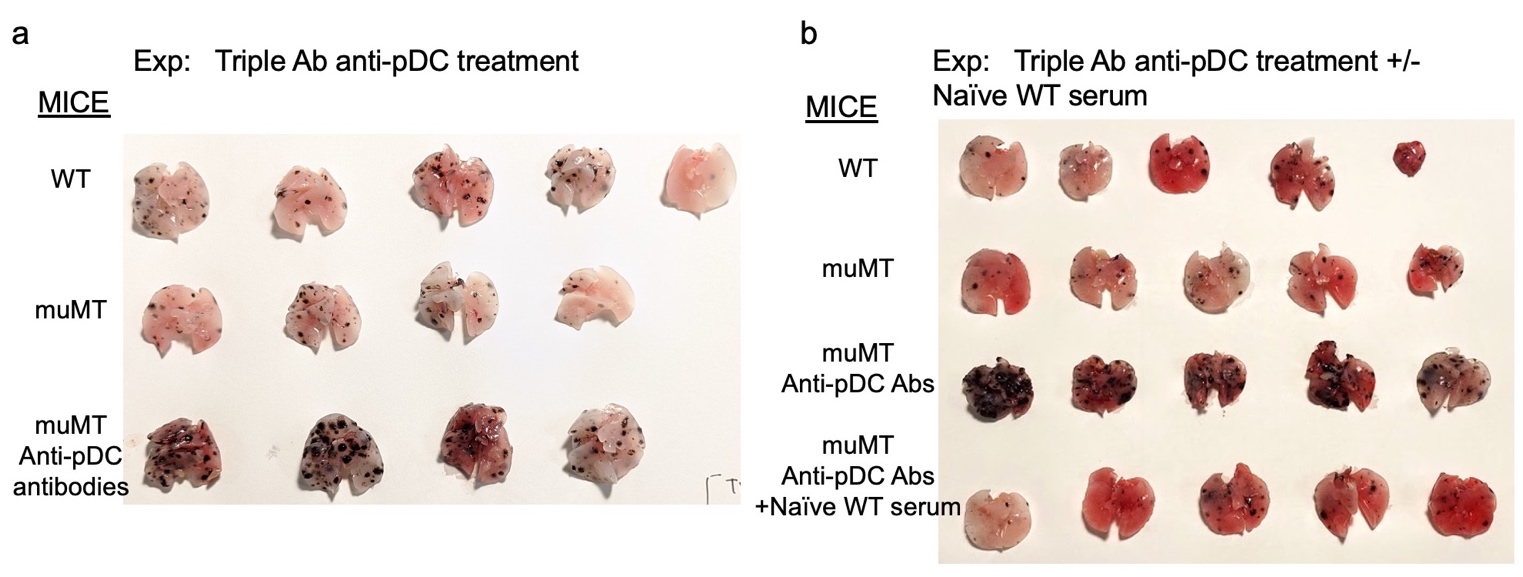

**Extended Data Fig. 5**| **Anti-pDC treatment alone increased the tumor burden in muMT mice, but anti-pDC treatment given with naive WT serum reduces the tumor burden.** Picture depicts agarose inflated lungs challenged with B16F10 melanoma in (a) WT, muMT, and pDC-depleted muMT; and (b) WT, muMT, pDC-depleted muMT, and pDC-depleted muMT injected with naive WT serum.
